# Cell assemblies in the cortico-hippocampal-reuniens network during slow oscillations

**DOI:** 10.1101/474973

**Authors:** David Angulo-Garcia, Maëva Ferraris, Antoine Ghestem, Lauriane Nallet-Khosrofian, Christophe Bernard, Pascale P Quilichini

## Abstract

The nucleus reuniens (NR) is an important anatomical and functional relay between the medial prefrontal cortex (mPFC) and the hippocampus (HPC). Whether the NR controls neuronal assemblies, a hallmark of information exchange between the HPC and mPFC for memory transfer/consolidation, is not known. Using simultaneous LFP and unit recordings in NR, HPC and mPFC in rats during slow oscillations under anesthesia, we identified a reliable sequential activation of NR neurons at the beginning of UP states, which preceded mPFC ones. NR sequences were spatially organized, from dorsal to ventral NR. Chemical inactivation of the NR disrupted mPFC sequences at the onset of UP states as well as HPC sequences present during sharp-wave ripples. We conclude that the NR contributes to the coordination and stabilization of mPFC and HPC neuronal sequences during slow oscillations, possibly via the early activation of its own sequences.

**Significance Statement:** Neuronal assemblies are believed to be instrumental to code/encode/store information. They can be recorded in different brain regions, suggesting that widely distributed networks of networks are involved in such information processing. The prefrontal cortex, the hippocampus and the thalamic nucleus reuniens constitute a typical example of a complex network involved in memory consolidation. In this study, we show that spatially organized cells assemblies are recruited in the nucleus reuniens at the UP state onset during slow oscillations. Nucleus reuniens activity appears to be necessary to the stability of prefrontal cortex and hippocampal cell assembly formation during slow oscillations. This result further highlights the role of the Nucleus Reuniens as a functional hub for exchanging and processing memories.

## Introduction

Information exchange between the hippocampus (HPC) and the medial prefrontal cortex (mPFC) is essential for different memory processes, including consolidation (Siapas and Wilson, 1998; Frankland and Bontempi, 2005; Preston and Eichenbaum, 2013; Maingret et al., 2016; Eichenbaum, 2017; Kitamura et al., 2017; Latchoumane et al., 2017; Ferraris et al., 2018). In both regions, the representation of information is supported by the recruitment of cell assemblies of neurons, which fire in a fine time-resolved manner (Skaggs and McNaughton, 1996; Nadasdy et al., 1999; Lee and Wilson, 2002; Euston et al., 2007; Luczak et al., 2007; Pastalkova et al., 2008; Peyrache et al., 2009; Battaglia et al., 2011). These neuronal assemblies display a precise temporal sequential activation, which reflects the encoded information (Nádasdy, 2000; Foster and Wilson, 2006; Ji and Wilson, 2007; Davidson et al., 2009; Marre et al., 2009; Pfeiffer, 2017). Such activity occurs mainly during sleep (Foster, 2017), particularly during non-REM sleep, and is organized in space and time by a set of oscillations, such as hippocampal sharp-wave ripples, cortical slow oscillations and spindles (Sirota et al., 2003; Sirota and Buzsáki, 2005; Staresina et al., 2015; Maingret et al., 2016). How these cells assemblies are finely coordinated between the HPC and the mPFC to support memory consolidation is not well understood. The thalamic nucleus reuniens (NR), which bi-directionally connects the HPC and the mPFC (Herkenham, 1978; Van der Werf et al., 2002; Vertes, 2006; Varela et al., 2014), plays a key role in memory consolidation (Loureiro et al., 2012; Cassel et al., 2013; Pereira de Vasconcelos and Cassel, 2015; Dolleman-van der Weel et al., 2019). In particular, the NR synchronizes gamma bursts, which can open temporal windows for information exchange (Buzsáki and Draguhn, 2004), between the HPC and the mPFC during non-REM sleep (Ferraris et al., 2018). The NR is therefore ideally poised to orchestrate the dynamics of cell assemblies in both regions.

We have recorded and manipulated the NR–mPFC–HPC circuit during slow oscillations under anesthesia, which resembles sleep patterns and provide long periods for analysis. We find that cell assemblies are recruited in the NR at the onset of the UP state of slow oscillations and that NR activity is needed for the stability of HPC and mPFC cell assemblies.

## Materials and Methods

### Contact for Reagent and Resource Sharing

Further information and requests for resources may be directed to and will be fulfilled by the Lead Contact, Dr. Pascale P. Quilichini.

### Experimental Model and Subject Details

All experiments were performed in accordance with experimental guidelines approved by Aix-Marseille University Animal Care and Use Committee. All rats were group housed to avoid social isolation-induced stress (Manouze et al., 2019). A total of 18 rats were used in this study. Part of these data (14 Wistar Han rat data) was used in a previously published study (Ferraris et al., 2018), and 4 Wistar Han rats are original data. They include local field potentials (LFPs) and single-unit recordings made in the mPFC, HPC and NR of anesthetized rats.

### Animal surgery

Wistar Han IGS male rats (250-400g; Charles River; RRID:RGD_2308816) were anesthetized with urethane (1.5 g/kg, i.p., Cat#U2500, Sigma-Aldrich) and ketamine/xylazine (20 and 2 mg/kg, i.m., Renaudin Cip: 3400957854195 and CENTRAVET Cat#ROM001, respectively), additional doses of ketamine/xylazine (2 and 0.2 mg/kg) being supplemented during the electrophysiological recordings. The heart rate, breathing rate, pulse distension and the arterial oxygen saturation were also monitored with an oximeter (MouseOx, Starr Life Science) during the entire duration of the experiment to ensure the stability of the anesthesia and monitor the vital constants. The head was secured in a stereotaxic frame (Kopf #962, Phymep) and the skull was exposed and cleaned. Two miniature stainless-steel screws, driven into the skull, served as ground and reference electrodes. Up to three craniotomies were performed to target, from bregma: the prelimbic area of the medial prefrontal cortex (mPFC) at +3 mm AP and +0.8 mm ML; the CA1 field of the intermediate hippocampus (HPC) at −5.6 mm AP and +4.3 mm ML; and the nucleus reuniens (NR) at −1.8 mm AP and −2 mm ML. Silicon probes (NeuroNexus) were used to record from these structures: a A1×32-Edge-5mm-20-177-H32-15 probe placed at [−2.5 - 3.1] mm from brain surface to reach mPFC layer 5; a A1×32-Edge-10mm-20-177-H32-50 32-site probes placed at −7.2 mm from brain surface to reach the NR; a HPC A1×32-6mm-50-177-H32-15 probe placed at [−2.8 −3.0] mm perpendicularly to the CA1 field from *stratum oriens* to *stratum lacunosum moleculare* in the HPC. All the probes were lowered inside the brain with a motorized manipulator (IVM single, Scientifica).

For NR inactivation experiments (n = 5 rats), a local injection of a fluorophore-conjugated muscimol (BODIPY-MSCI TMR-X Conjugate, Cat#M23400, Invitrogen) was performed in the NR and data from the mPFC and the HPC (CA1) were simultaneously acquired. The injection needle (33 gauge, Cat#87930, Cat#7803-05, Hamilton) was inserted in the NR (using the same depth coordinates as the probes and mounted on the same micromanipulator) and 0.70 nmol of muscimol in 0.3 μl of PBS (Ferraris et al., 2018) was delivered over 60s through a micropump (UltraMicroPump UMP3-1, WPI). The needle was left in place for 3 additional minutes to allow for adequate diffusion of the drug, then carefully removed.

At the end of the recording, the animals were injected with a lethal dose of Pentobarbital Na (150mk/kg, i.p.) and perfused intracardially with 4% paraformaldehyde solution in phosphate buffer (0.12M). The position of the electrodes (DiIC18(3), Cat#46804A, InterChim) was applied on the back of the probe before insertion) was confirmed histologically on Nissl-stained 60 μm sections (NeuroTrace 500/5225 Green Fluorescent Nissl Stain, Cat#N21480, Invitrogen). Only experiments with appropriate position of the probe were used for analysis (Fig. 2A).

### Data collection and initial analysis

Extracellular signal recorded from the silicon probes was amplified (1000x), bandpass filtered (1 Hz to 5 kHz) and acquired continuously at 32 kHz (64-channel DigitalLynx; NeuraLynx) at 16-bit resolution. Raw data was preprocessed using a custom-developed suite of programs (Csicsvari et al., 1999). After recording, the signals were downsampled to 1250 Hz for the LFP. Spike sorting was performed automatically, using KLUSTAKWIK ((Harris et al., 2000); http://klustakwik.sourceforge.net/; RRID:SCR_008020; RRID:SCR_014480), followed by manual adjustment of the clusters, with the help of autocorrelogram, cross-correlogram and spike waveform similarity matrix (KLUSTERS software, RRID:SCR_008020; (Hazan et al., 2006). After spike sorting, the spike features of units were plotted as a function of time, and the units with signs of significant drift over the period of recording were discarded. Moreover, only units with clear refractory periods and well-defined cluster were included in the analyses. Recording sessions were divided into brain states of theta and slow oscillation periods. The epochs of stable slow oscillations (SO) periods were visually selected respectively from the ratios of the whitened power in the slow oscillations band (0.5-2 Hz) and the power of the neighboring band (20-30 Hz) of mPFC or NR LFP and assisted by visual inspection of the raw traces (Quilichini et al., 2010).

Neurons were assigned as “NR neurons” by determining the approximate location of their somata relative to the recording sites, the known distances between the recording sites, the histological reconstruction of the recording electrode tracks and subsequent estimation of the recording sites. All the neurons recorded from sites located near the close contour of the NR were discarded. Neurons located at a minimal distance of 200μm of NR border and located within contours of the ventro-median, submedian or antero-median thalamic nuclei were classified as “other thalamic neurons” and used in the analysis (Ferraris et al., 2018).

### Data post-processing

All the analysis was performed using custom-written MATLAB (RRID:SCR_001622; RRID: SCR_008020) scripts. From the spike times, the instantaneous firing rates of each cell was calculated by counting the number of spikes inside a window of 50 ms, in overlapping intervals of 10 ms. The population firing rate was estimated averaging the single cell firing rates at each 10ms interval. Both the population rate and the single cell firing rate were smoothed with a Gaussian kernel of 5ms width. Peaks of population activity (AP) were identified as the points where the population rate amplitude was larger than the average rate plus 1 standard deviation. A separation between two consecutive peaks of at least 600 ms was imposed on the peak detection algorithm to avoid multiple peaks of activity within one up-state. For visualization purposes, all firing rate’s heat maps were normalized with the peak firing rate for each cell to guarantee a variation between [0:1].

### Phase analysis and sequence identification

LFP signals (from mPFC or NR) during slow oscillation phase were band-pass filtered between 0.5 Hz and 2 Hz with a second order Butterworth filter to extract only the UP/DOWN transitions. The time evolution of the phase during the UP/DOWN cycle was extracted performing the Hilbert transform of the filtered LFP (from the mPFC or the NR, where the [0.5 2] Hz oscillation is most likely volume-conducted from the cortex). We used Rayleigh circular statistics (Berens, 2009; Ferraris et al., 2018) to compute the mean phase at which each neuron fires (“preferred phase”) and to build their firing-phase histograms (Fig. 1Ba). For visualization purposes, the resulting histograms depicted in the heat maps were normalized with the peak value of the distribution. For each cell, we then calculated a resultant vector characterized by an angle describing the average preferred phase and a magnitude with values between [0 1], quantifying the coherence of the phases. The template order was obtained by organizing the average preferred phase in increasing order (0 to 2π).

**Figure 1.**
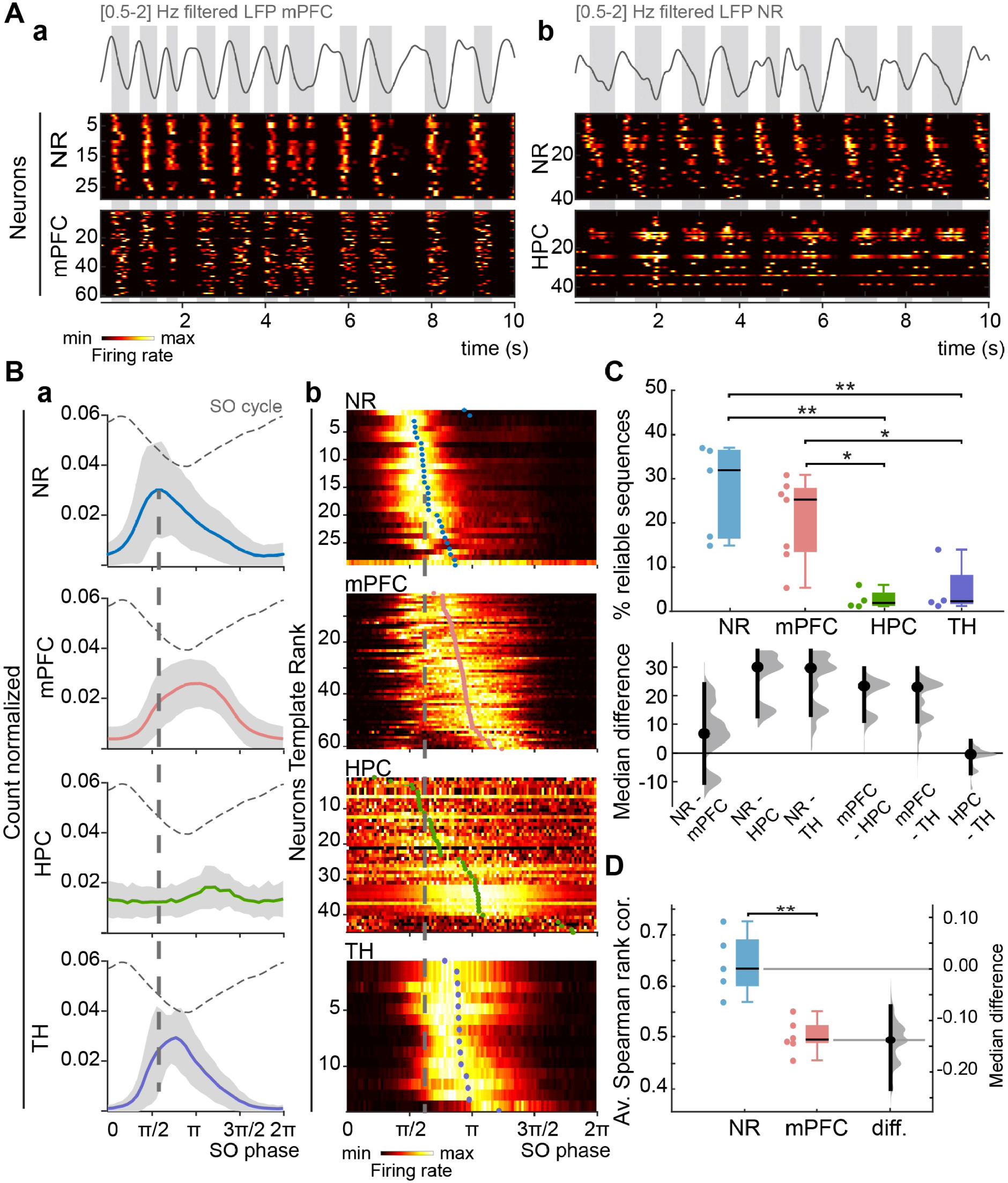
Sequential dynamics of cell assemblies. **A)** The upper trace depicts the [0.5-2] Hz filtered LFP in **(a)** mPFC and **(b)** NR (which is volume-conducted from the cortex), where the UP states correspond to troughs in the LFP signal (gray boxes), from two typical recordings. The corresponding template heat maps (bottom) of normalized NR unit activities with **(a)** mPFC or **(b)** HPC normalized unit activities show the repetition of neuronal assemblies particularly in the NR and the mPFC. The identification number of the recorded neurons are indicated on the left axis of the heat-maps. **B) (a)** Phase distribution of the NR, mPFC, HPC and TH population firing (grouped data, all recordings) with reference to the SO phase (depicted by the dashed curve). The dashed line indicates the most preferred phase of NR neurons, showing that their firing precedes other cells located in the mPFC, HPC and TH at the onset of the UP state. **(b)** Heat maps of normalized distribution of neurons preferred SO phase for sample representative firing patterns of NR (same neurons as in Aa), mPFC (same as in Aa), HPC (same as in Ab) and TH (representative example). The activation order for each neuron is calculated as the average preferred phase. Heat maps are ordered according to the increasing value of average preferred phase (colored circles), defining a template order. The sequential activation is clearly apparent in NR and mPFC neurons, but neither in the HPC nor in the TH (the phase distribution extends over at least p/2 for each neuron). **C)** Top panel: The largest percentage of reliable sequences is found in the NR and mPFC as compared to the HPC and the TH. Bottom panel: the median differences for 6 comparisons are shown in the above Cumming estimation plot. Each median difference is plotted on the lower axes as a bootstrap sampling distribution. Median differences are depicted as dots; 95% confidence intervals are indicated by the ends of the vertical error bars. **D)** The average Spearman correlation shows that there are more reliable sequences in the NR than in the mPFC. The estimation plot on the right side shows the median difference (dots) and the 95% confidence interval.

**Figure 2.**
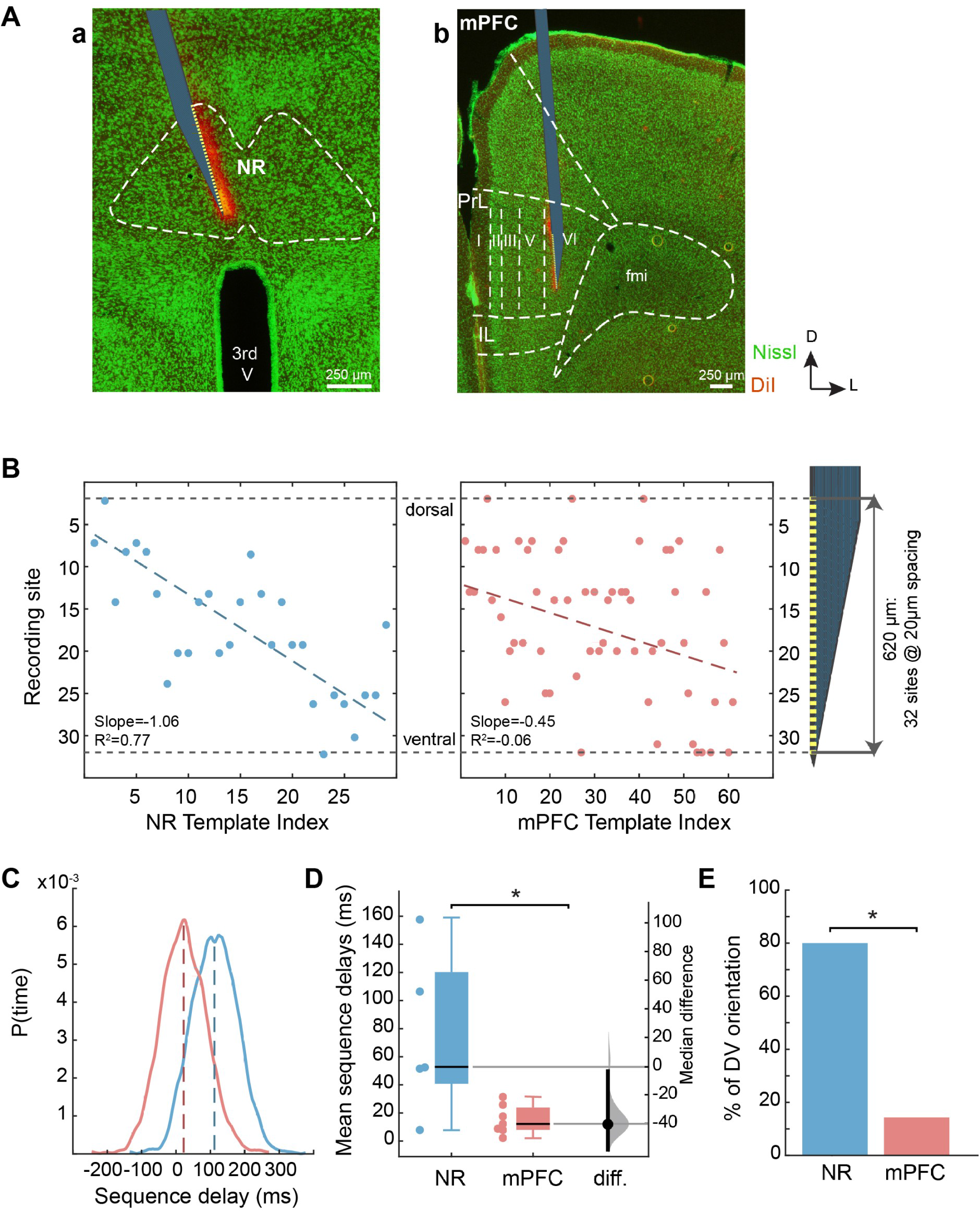
Spatial organization of NR and mPFC cell assemblies. **A)** Representation of the position of a linear silicon probe with 32 recording sites in **(a)** the NR and **(b)** mPFC. NR and mPFC contours (and layers) are delimited by the white dashed line over the green fluorescent Nissl staining (fmi: forceps minor of the *corpus callosum*; 3^rd^ V: third ventricle; PrL: prelimbic area; IL: infralimbic area). The red-orange staining corresponds to the DiI that was deposited at the back of the silicon probe before insertion (D: dorsal; L: lateral). **B)** Relationship between anatomical dorso-ventral location of NR (left panel) and mPFC (right panel) neurons (location defined by the site of the probe recording the maximum amplitude of the action potentials) and the template rank showing a linear correlation for NR neurons but not for mPFC ones in a template experiment. **C)** Distribution of spatial delays (from the beginning to end of a sequence) across the probe from the topmost site to the bottom-most one for mPFC (red) and NR (blue) neurons. **D)** Mean sequence delays (with median difference and 95% confidence interval) across experiments for NR and mPFC. **E)** The percentage of experiments showing a dorso-ventral (DV) orientation of sequences was larger in the NR than in the mPFC.

### Sequence reliability, participation index and sequence delay

We extracted the UP state duration as described in Ferraris et al. (2018). At each UP state, the local activation order was calculated by measuring the time lag between the first activation of each neuron relative to the AP in a +/− 200ms window. Ordering the latencies in increasing order resulted in the local activation order. The reliability of a given sequence within a population peak was quantified as the Spearman rank correlation of the template order and the local activation order in that particular UP state. Sequences were considered reliable above 99% significance level. For each reliable sequence, a robust fit between the local activation order and the activation latency was computed. To assess whether a neuron participated in a given sequence, the outliers in the robust fit were identified as those whose residual value were larger than twice the standard deviation of the residuals in the robust fit. The participation index was then calculated as the mean fraction of neurons that participated in the sequences of a recording session. A second linear fit between the local activation latency and the site of the linear probe closest to the neuron gave a measure of the directionality of the sequence activation. A slope of the fit different than 0 implies an activation towards a given direction. Since this slope has units of time/electrode site, we multiplied the resulting value by 32 (the number of sites in the electrode) to account for the time required for a sequence (sequence delay) to travel along the electrode.

### Activity peak triggered histogram (APTH)

Once the APs have been identified according to the algorithm described in Data post-processing section, we store the values of activation time lag relative to the AP at each UP state, allowing us to compute the so-called activity peak triggered histogram for each cell. With the APTH it is possible to assess the statistics of the firing latencies in the neighborhood of the UP state (+/− 200 ms). Once the APTH was obtained, one can calculate the variability of the activation lags as a measure of the firing extent around the peak reported in Fig. 3C. Variability of the APTH can be calculated in analogy to the variance of a probability distribution function where τ is the mean value of the APTH (See Fig. 3C insert).

**Figure 3.**
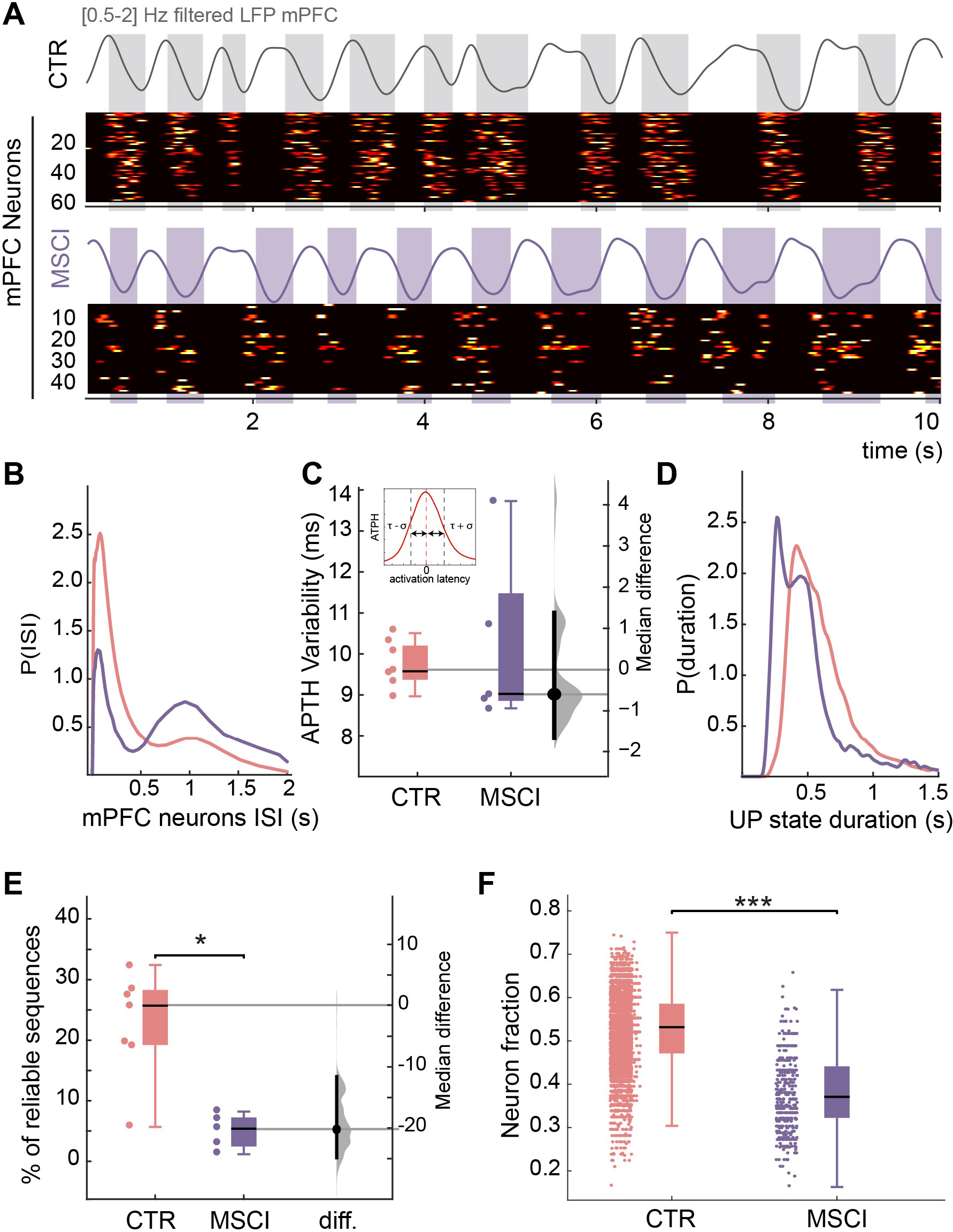
Inactivation of NR impairs reliable sequential activation of mPFC neurons at the beginning of the UP state. **A)** Two template heat maps of normalized mPFC unit activities in control and during NR inactivation condition showing less stable cell assemblies. The upper traces depict the [0.5-2] Hz filtered LFP in mPFC, where the UP states correspond to troughs in the LFP signal. **B)** Inter-spike interval distribution of mPFC neurons grouped data for control (red) and NR inactivation conditions (purple), showing that a decrease in burst firing when the NR is inactivated. **C)** Grouped data with median difference and 95% confidence interval of the variability of the APTH for control (CTR) and NR inactivation conditions (MSCI). Insert: APTH: the activity peak triggered histogram quantifies the lag distribution with respect to the population activity peak for each neuron. **D)** Distribution of the duration of the UP state in control (red) and NR inactivation (purple) condition, where a peak at short values indicates the emergence of shorter UP states. **E)** The number of reliable sequences found in mPFC was decreased when the NR was inactivated (MSCI) as compared to the control (CTR) condition (grouped data, n=5 experiments). **F)** The fraction of neurons participating to reliable sequences was decreased when the NR was inactivated.

### SPW-Rs detection and analysis

The procedure of SPW-Rs detection in the HPC stratum pyramidale LFP was based on those described previously (Isomura et al., 2006; Ferraris et al., 2018). Briefly, the LFP was digitally bandpass filtered [80 250] Hz, and the power (root-mean-square) of the filtered signal was calculated. The mean and SD of the power signal were calculated to determine the detection threshold. Oscillatory epochs with a power of 5 or more SD above the mean were detected. The beginning and the end of oscillatory epochs were marked at points where the power fell 0.5 SD. Once the SPW-Rs were detected, the SPW-R half time was calculated as the average between the start and the end of it. Then, the APTH for each cell was computed, taking each SPW-R half time as the activity peak. To test whether a cell robustly participates in the SPW-Rs, we compared the APTH against a uniform distribution with identical mean and standard deviation. Cells whose APTH were different from the flat distribution above a 95% level were considered as robustly participating in the SPW-Rs. Cell participation ratio was obtained dividing the number of robustly participating neurons by the total number of recorded neurons for that session. Organizing the average time lag for robustly participating neurons in increasing order defined the template order for the ripple. For each SPW-R, the reliability of activation respect to the template was calculated via the Spearman rank correlation between the activation order of that SPW-R and the template order. To calculate this correlation, only the neurons that belong to the template order were considered. A correlation above 95% confidence interval was considered reliable.

### Statistics

All results reported are based on a significance threshold α=0.05, otherwise stated, and all groups included enough samples to enable rejection of the null hypothesis at that level. We used two sample Kolmogorov-Smirnov test to assess differences between distributions, and t-Student’s test to evaluate differences in the mean of distributions. In parallel, results from small samples data (n < 10) were evaluated using estimation statistics providing the effect size and the 95 % confidence interval (CI) of the median difference between two groups. Raw data was processed online in https://www.estimationstats.com website directly giving both results and graphs (Calin-Jageman and Cumming, 2019). The median difference between two groups is shown with a Gardner-Altman estimation plot. Raw data is plotted on the left axis. The median difference is plotted on a floating axis on the right as a bootstrap sampling distribution (Fig. 1CD 2D, 3CE, 4B). Five thousand bootstrap samples were taken; the confidence interval is bias-corrected and accelerated. The mean difference is depicted as a dot; the 95 % confidence interval is indicated by the ends of the vertical error bar. If 95% CI includes 0, differences are considered as non-significant. The p value(s) reported are the likelihood(s) of observing the effect size(s), if the null hypothesis of zero difference is true. Correlation tests involving ranked variables (neuron indices and electrode sites) were performed via a Spearman rank correlation. We tested significant differences between percentages with a two proportion Z-test.

**Figure 4:**
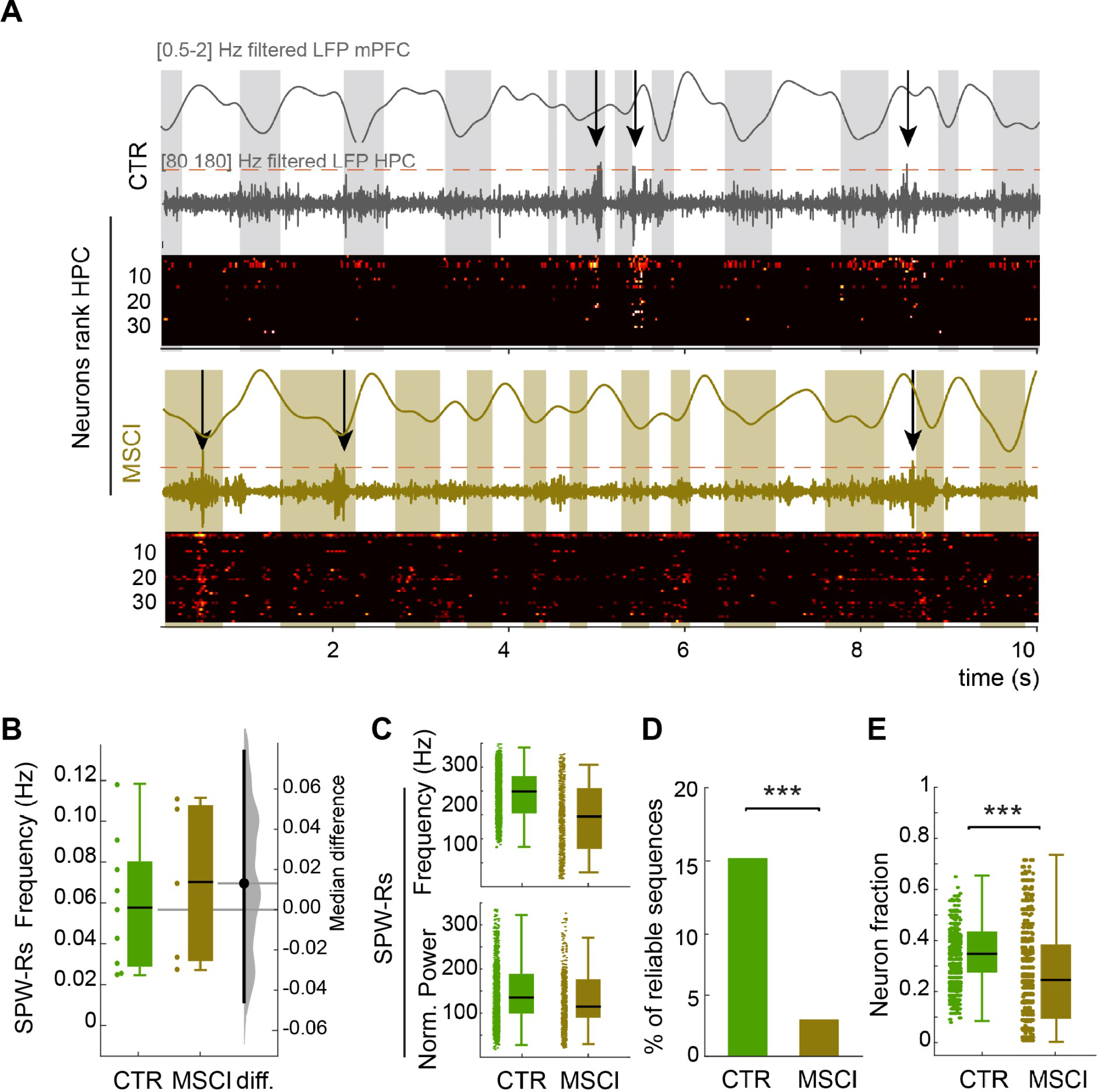
Inactivation of NR impairs reliable sequential activation of HPC neurons during SPW-Rs. **A)** Two template heat maps of normalized HPC unit activities in control (CTR) and NR inactivation (MSCI) condition(bottom of each panel). The upper traces depict the [0.5-2] Hz filtered LFP in mPFC, where the UP states correspond to troughs in the LFP signal, and the black arrow mark the presence of a SPW-R event, which are visible in the [80 200] Hz filtered LFP (bottom trace). **B)** SPW-Rs occurrence did not changed when NR was inactivated (grouped data, n=5 experiments, n=913 SPW-R events in CTR, n=149 SPW-R events in MSCI). **C)** NR inactivation did not change the normalized power and inner frequency of SPW-Rs. **D)** The proportion of reliable cell assemblies during SPW-Rs was considerably reduced when the NR was inactivated. **E)** The fraction of reliably participating HPC neurons to sequences during SPW-R events was decreased in NR inactivation condition.

## Results

### Cell assemblies are recruited in NR at UP state onset

We first assessed the behavior of NR (n=166 cells, n=5 rats), mPFC (n=496, n=7) and HPC (n=163, n=4) neurons during slow oscillations (SO) in anesthetized rats. Slow oscillations during anesthesia share similar features as non-REM sleep (Hutt, 2011; Ferraris et al., 2018) and offer the advantage of long duration stable recordings necessary to identify statistically significant sequences. Neighboring thalamic neurons (antero-median and ventro-median nuclei) were also recorded as a control group (“TH”, 89 neurons, n=4).

During SO, most of the neurons fired at the onset of the UP states, mainly in mPFC and NR (Fig. 1A) (Ferraris et al., 2018). We found a robust recruitment of neuronal activity at the UP state onset in NR (median percentage of recruited neurons per UP state 70.4 %, min/max: 61.1 / 85.9 %), and mPFC (median: 62.9 %, min/max: 42.1 / 70 %). In comparison, HPC neurons showed less participation during UP state (median: 33.8 %, min/max: 29.3 / 46.4 %. To identify any ordering in this firing activity, we ranked each neuron according to its mean preferred SO phase (Fig. 1Ba). In contrast, HPC neurons displayed better sequential activation during sharp-wave ripples (SPW-Rs), which tend to occur during the UP state (Sirota et al., 2003; Battaglia et al., 2004; Isomura et al., 2006; Buzsáki, 2015; Maingret et al., 2016; Khodagholy et al., 2017) (median: 45 %, min/max: 35 / 64 %). Neurons were organized in increasing order of average preferred phase, defining a *template order* (Fig. 1Bb). Sequences in NR always preceded mPFC ones (Fig. 1B). As a negative control, we analyzed the activity of neighboring thalamic nuclei (TH) neurons. Although TH neurons displayed a strong entrainment by SO (Ferraris et al., 2018), there was poor sequential activity (Fig. 1B). The sequential activity at UP state onset therefore appears to be specific of NR and mPFC neurons in the areas investigated here. As a further control, we analyzed the activity of NR neurons during epochs dominated by theta (4-6 Hz) oscillations. We did not find stable sequential neuronal assemblies during theta (data not shown). Together, these results show that, at the beginning of UP states, cell assembly formation occurs first in the NR, then in the mPFC and marginally in the HPC and neighboring thalamic nuclei.

We then quantified the reliability of these sequences. To do so, we computed at each UP state the activation latency of each neuron relative to the population peak activity (see Methods). The local activation order of the cell assembly was determined from the ascending sorting of the activation latencies. We then calculated the Spearman rank correlation between the local activation order and the template order. A rank correlation r = 1 indicates that the sequential activation follows exactly the template order determined previously by the average phase preferences. For NR neurons, the template order was significantly expressed in 32% of the total number of UP states (min/max = 15 / 37 %, Spearman test, p < 0.01), while mPFC reliable sequences were found in 25 % of the UP states (min/max = 5 / 31 %, Fig. 1C). In contrast, reliable HPC sequences were only detected in 2 % of the UP states (min/max = 1 / 6 %) and in 2 % for sequences in TH neurons (min/max = 1 / 14 %). Although the proportion of reliable sequences in the NR and mPFC were not significantly different (KS-test, p = 0.15), the average Spearman correlation values of the reliable sequences were significantly larger in the NR than in the mPFC (KS-test, p < 0.01, Fig. 1D), indicating that the NR cell assembly sequential organization is the most consistent. To estimate the participation of neurons in sequence generation, we calculated a participation index, defined as the probability for a given neuron to be involved in a given sequence (see Methods). The participation index was consistently larger across experiments in the NR (median: 0.63, min/max: 0.55 / 0.81) than in the mPFC (median: 0.52, min/max: 0.36 / 0.60; KS-test, p < 0.05). The sequential firing of neurons could just be the result of an increased excitability of early firing neurons as compared to neurons firing later after the onset of the UP state (Buzsáki, 1989). We thus tested the correlation between the firing order and the firing rate for each NR and mPFC neuron. We found that 2/5 NR recordings were significantly correlated while only 1/7 mPFC recordings showed significant correlations (Spearman test, p<0.05).

Together, these results show that mPFC and NR neurons display highly stable patterns of sequential neuronal activations at the onset of the UP state, with NR sequences being more reproducible. The sequential activation cannot be solely explained by increased excitability of early firing cells.

### NR cell assemblies are spatially distributed

We also investigated the spatial distribution of the NR and mPFC cell assemblies. To do this, we correlated the template order of each neuron to their anatomical dorso-ventral localization, since the silicon probes was sampling NR from the dorsal to ventral pole (Fig. 2A). The slope of a linear fit of such correlation provides the information on whether the activity is propagating in a given direction (Luczak et al., 2007) (see Methods, Fig. 2B). Furthermore, we quantified the degree of spatial distribution of the activity for each UP state, calculating a linear fit between the activation latency of each neuron relative to the peak activity and the dorso-vental localization. The slopes of the fit for each UP state, properly multiplied by the number of sites of the probe resulted in the sequence delay (see methods). Positive or negative delays are fingerprints of propagating activity in certain direction. Even though both NR and mPFC can show some degree of propagation in a preferential direction (Fig. 2C), NR average delays were consistently larger than mPFC ones (KS-test, p < 0.05, Fig. 2D), reflecting a dorsal-to-ventral direction of the sequential activation. To assess the consistency of the preferential direction of propagation, we calculated the Spearman rank correlation between the rank of the template order and the anatomical location of the corresponding recording site. We found a highly significant correlation in NR recordings in most cases (n = 4 out of 5, median | r | value across all n: 0.65, p < 0.001), whereas it was virtually absent in mPFC (n = 1 out of 7, |r| = 0.35, p < 0.01). There are thus more spatially organized sequences in NR as compared to mPFC (z-test, p < 0.05, Fig. 2E).

Altogether, these results demonstrate that NR displays at the UP state onset robust sequential activations, which are spatially organized in a dorso-ventral pattern. This is particularly surprising as it suggests a topical spatial organization in the NR.

### NR activity is necessary to the mPFC and HPC sequences stability

The fact that NR cell assemblies occur earlier than mPFC ones raises the possibility that the NR may be involved in the sequential activation of mPFC neurons. To test this hypothesis, we used Muscimol to inactivate chemically the NR (see Methods) and recorded the activity of mPFC neurons (n=190 neurons, n=5 experiments). Following NR inactivation, mPFC neurons showed a reduced firing activity during the UP state as compared to non-inactivation, control recordings (Fig. 3A) (Ferraris et al., 2018). During SO, the distribution of interspike intervals is characterized by the presence of two peaks: one around 1 Hz corresponding to the frequency of the SO, and one around 100 ms, which indicates that cells fire bursts of action potentials (Fig. 3B). The distribution of interspike intervals was modified in inactivation conditions with a smaller peak around 100 ms, reflecting reduced bursting activity during UP state (Fig. 3B). We then calculated the distribution of the pooled variability of the activity peak triggered histogram (APTH, see methods and Fig. 3C insert) in control and NR inactivation conditions. The median APTH variability across experiments was not significantly changed when NR was inactivated (control median: 9.62, min/max: 8.98 / 10.60, NR inactivation median: 9.02, min/max: 8.67 / 13.73, KS-test, p =0.4428), as shown in Fig. 3C. This reveals the fact that, even though bursting is reduced, the time extent to which neurons in mPFC fire is maintained. However, the UP state duration was slightly shorter (control: 0.55 s, NR inactivation: 0.48 s, Mann-Whitney test, p < 0.001), an effect that can be attributed to the extra peak in UP state duration distribution around 0.25 s, which did not exist in control data (Fig. 3D). We then evaluated the outcome of the mPFC sequences in the NR inactivation condition. Inhibition of NR activity reduced the capacity of mPFC to generate reliable sequences as compared to the control condition, since only 5.6% of them (min/max: 1.4 / 8.4%) were reliable (as compared to control median: 25%, min/max: 5 / 30 %, KS-test, p=0.0228 Fig. 3E). Moreover, the fraction of participating neurons in reliable sequences decreased in the inactivation condition (control: 53%; inactivation: 37%; KS-test, p < 1e-5; Fig. 3F). These results support the proposal that NR activity is important for the stability and reliability of mPFC sequences.

The UP state of the SO poorly modulates the activity of HPC neurons and the formation of timely organized cell assemblies. However, sequential activation of HPC neurons consistently occurs during SPW-Rs (Buzsáki, 2015). We therefore investigated the consequences of NR inactivation on HPC sequences during SPW-Rs. NR inactivation did not affect HPC neurons firing rate (control median: 70 Hz, min/max: 41 / 85 Hz, NR inactivation median: 52.59 Hz, min/max: 33 / 71 Hz, KS-test, p=0.97, Fig. 4A). In addition, NR inactivation did not alter the frequency of SPW-Rs occurrence (median control: 0.087 Hz, min/max: 0.043 / 0.11 Hz, NR inactivation: 0.049 Hz, min/max: 0.022 / 0.111 Hz, KS-test, p=0.8254, Fig. 4B). Similarly, neither their power (mean normalized power control: 157 ± 2, NR inactivation: 144 ± 5, T-test, p=0.295; Fig. 4C bottom panel) nor their inner frequency (mean frequency control: 193 ± 1 Hz, NR inactivation: 213 ± 3 Hz, T-test, p=0.084, Fig. 4C top panel) were modified. This result shows that NR inactivation does not change either SPW-R properties or global HPC cell firing. In contrast, the number of reliable sequences found within SWP-Rs was drastically reduced when the NR was inactivated (control median: 15%, NR inactivation median: 3%, z-test, p<0.001; Fig. 4D). In this condition, the remaining cell assemblies recruited a significantly lower number of neurons (control median: 37.5 %, min/max: 12.5% / 80%, NR inactivation median: 27.8 %, min/max: 0% / 74.4%, KS-test, p<0.001, Fig. 4E). These findings suggest that the NR also controls the sequential organization of neuronal firing in the HPC during SWP-Rs.

## Discussion

In this study, we show that two third of NR neurons fire within robustly spatially and temporally organized cell assemblies at UP state onset; and that NR activity controls cell assemblies’ stability in mPFC at UP state onset and in HPC during SPW-Rs. These results further support the concept that the NR is a key functional hub in memory networks involving the medial prefrontal cortex and the hippocampus.

The sequential activation of neuronal assemblies is believed to constitute a core feature of information processing in the brain (Tonegawa et al., 2018). Cell assemblies are found in the archicortex (Lee and Wilson, 2002; Harris et al., 2003; Dragoi and Buzsáki, 2006; Pastalkova et al., 2008; Villette et al., 2015; Malvache et al., 2016), cortical areas (Kenet et al., 2003; MacLean et al., 2005; Ferezou et al., 2006; Euston et al., 2007; Luczak et al., 2007; Luczak and Maclean, 2012), as well as in striatum (Lansink et al., 2009). They constitute a way to code/encode/store information (Maass, 2016; Kitamura et al., 2017). During non-REM sleep, cortical activity is dominated by the sequential activation of cortical neurons at the onset of the UP state, while, in the hippocampus, the sequential activation mostly occurs during SPW-Rs (Sirota et al., 2003; Battaglia et al., 2004; Peyrache et al., 2009; Maingret et al., 2016; Khodagholy et al., 2017). Similar sequential firing occurs during the slow oscillations measured during anesthesia (Luczak et al., 2007), supporting the view that such brain state shares many features with non-REM sleep (Tung and Mendelson, 2004; Isomura et al., 2006; Clement et al., 2008; Quilichini et al., 2010; Hutt, 2011; Ferraris et al., 2018). Whether sequences represent internally generated representations or preconfigured cell assemblies (Pastalkova et al., 2008; Dragoi and Tonegawa, 2012; Liu et al., 2019) or a functional template of offline replay in the framework of memory consolidation (Lee and Wilson, 2002; Buzsáki, 2015; Pfeiffer, 2017) still remains to be elucidated. Our work demonstrates that sequences can also be recorded in a thalamic nucleus. This property appears to be specific to the NR, on the basis of recordings of other thalamic nuclei recorded in the vicinity. A systematic study of all thalamic nuclei should now be performed, to determine whether this property is specific to the NR. In the present work, we used anesthesia conditions, as the identification of sequences requires long lasting stable recordings (Bermudez Contreras et al., 2013). Our results should also be validated during long sleep sessions.

The way mPFC and HPC sequences are generated remains poorly understood. Our results demonstrate that NR activity is a key regulator of mPFC and HPC sequence stability. Interestingly previous (Ferraris et al., 2018) and present results demonstrate that the NR does not regulate oscillatory patterns, as NR inactivation does not change the frequency of SO, gamma oscillations and SPW-Rs. However, the NR seems to organize the synchronization of gamma oscillations between the HPC and the mPFC (Ferraris et al., 2018) as well as the sequential activation of cell assemblies in the mPFC during SO or in the HPC during SPW-Rs. The NR is ideally located for such fine-tuning of cell activity, as it is bi-directionally connected to the mPFC and HPC (Vertes, 2006; Cassel et al., 2013; Varela et al., 2014). The fact that NR sequences always precede mPFC ones at UP state onset suggests that NR cells may directly drive mPFC cells. In keeping with this proposal, NR neurons have an excitatory action on HPC and mPFC (Dolleman-Van der Weel et al., 1997; Dolleman-Van der Weel and Witter, 2000; Di Prisco and Vertes, 2006; Dolleman-van der Weel et al., 2019) by modulating/activating both interneurons and principal cells. However, the control of sequences during hippocampal SPW-Rs is more difficult to explain as SPW-Rs occur at variable times after UP state onset (Sirota et al., 2003; Battaglia et al., 2004). This rather suggests that the NR is an important hub in a wider network of networks, from which oscillations can emerge.

The presence of sequences specifically in the NR is quite remarkable. Since NR neurons are involved in reference memory consolidation (Loureiro et al., 2012) and in spatial memory (Jankowski et al., 2014; Ito et al., 2015; Jankowski et al., 2015; Ali et al., 2017; Cholvin et al., 2018), NR sequences may constitute an activity template used to organize information at the beginning of the UP state. Such a mechanism may enable the transmission of information to target areas in a packet-based manner (Luczak et al., 2015); a default activity in a default mode (Luczak and Maclean, 2012; Sanchez-Vives and Mattia, 2014).

Another remarkable feature of NR sequences is that their dorso-ventral organization, suggesting a precise topological organization in terms of afferences and efferences, despite the fact that NR does not have a layered organization (Jones, 1985; Bokor et al., 2002; Van der Werf et al., 2002). The NR probes was inserted in the brain with an angle to properly target the NR (Fig.. 2Aa). As a result, the bottommost sites were located more medially than the topmost ones. Although the medio-lateral distance was less that 250 μm between the top and bottom sites, we cannot fully rule out that the spatial organization we describe is medio-lateral rather than dorso-ventral (or both). The origin of the dorso-ventral organization of these sequences remains to be elucidated. While NR outputs to the subiculum are topographically organized along the dorso-ventral axis, no study has reported such precise topological organization of NR inputs (Dolleman-Van Der Weel and Witter, 1996). Only differences involving the rostro-caudal axis have been so far reported (Cassel et al., 2013), as well as a preferential targeting of NR fibers to prelimbic and infralimbic deep layers (5 and 6) and superficial layer 1 in mPFC (Berendse and Groenewegen, 1991; Vertes, 2004; Hoover and Vertes, 2012). The fact that we could not find any dorso-ventral pattern in mPFC sequences does not mean that there is none to be found as, given the spatial organization of the mPFC, we could only target one layer. There is also no available information on the local connectivity among NR neurons, except a caudal to rostral pathway (Dolleman-Van der Weel et al., 1997). The NR includes difference cell types, but the lack of specific molecular markers prevents, so far, a proper optogenetic investigation (Bokor et al., 2002; Walsh et al., 2017).

In conclusion, our results further support the concept that the NR plays a key role as an anatomical and functional hub between the mPFC and the HPC. The control it exerts on mPFC and HPC information packets suggests that it strongly participates in the organization of information in both regions but also in the transfer of information from the HPC to the mPFC. Its internal organization allows the genesis of information packet sequences, which may represent similar features as those coded in the mPFC and the HPC.

## Acknowledgments

This work was supported by grants from the FRM (Fondation Recherche Médicale FDT201805005246 to M. F.). D.A-G received support by the A*MIDEX grant (No. ANR-11-IDEX-0001-02). We thank Wesley Clawson for proofreading this manuscript.

## Notes

**Declaration of Interests**: The authors declare no competing interests

## REFERENCES

Ali M, Cholvin T, Muller MA, Cosquer B, Kelche C, Cassel J-C, Pereira de Vasconcelos A (2017) Environmental enrichment enhances systems-level consolidation of a spatial memory after lesions of the ventral midline thalamus. Neurobiology of learning and memory 141:108–123.

Battaglia FP, Sutherland GR, McNaughton BL (2004) Hippocampal sharp wave bursts coincide with neocortical “up-state” transitions. Learning & memory 11:697–704.

Battaglia FP, Benchenane K, Sirota A, Pennartz CM, Wiener SI (2011) The hippocampus: hub of brain network communication for memory. Trends Cogn Sci 15:310–318.

Berendse HW, Groenewegen HJ (1991) Restricted cortical termination fields of the midline and intralaminar thalamic nuclei in the rat. Neuroscience 42:73–102.

Berens P (2009) CircStat: A MATLAB Toolbox for Circular Statistics. Journal of Statistical Software 31:1–21.

Bermudez Contreras EJ, Schjetnan AGP, Muhammad A, Bartho P, McNaughton BL, Kolb B, Gruber AJ, Luczak A (2013) Formation and reverberation of sequential neural activity patterns evoked by sensory stimulation are enhanced during cortical desynchronization. Neuron 79:555–566.

Bokor H, Csáki A, Kocsis K, Kiss J (2002) Cellular architecture of the nucleus reuniens thalami and its putative aspartatergic/glutamatergic projection to the hippocampus and medial septum in the rat. The European journal of neuroscience 16:1227–1239.

Buzsáki G (1989) Two-stage model of memory trace formation: A role for “noisy” brain states. Neuroscience 31:551–570.

Buzsáki G, Draguhn A (2004) Neuronal oscillations in cortical networks. Science 304:1926–1929.

Buzsáki G (2015) Hippocampal sharp wave-ripple: A cognitive biomarker for episodic memory and planning. Hippocampus 25:1073–1188.

Calin-Jageman RJ, Cumming G (2019) Estimation for Better Inference in Neuroscience. eNeuro 6.

Cassel JC, Pereira de Vasconcelos A, Loureiro M, Cholvin T, Dalrymple-Alford JC, Vertes RP (2013) The reuniens and rhomboid nuclei: neuroanatomy, electrophysiological characteristics and behavioral implications. Progress in neurobiology 111:34–52.

Cholvin T, Hok V, Giorgi L, Chaillan FA, Poucet B (2018) Ventral Midline Thalamus Is Necessary for Hippocampal Place Field Stability and Cell Firing Modulation. J Neurosci 38:158–172.

Clement Ea, Richard A, Thwaites M, Ailon J, Peters S, Dickson CT (2008) Cyclic and sleep-like spontaneous alternations of brain state under urethane anaesthesia. PloS one 3:e2004–e2004.

Csicsvari J, Hirase H, Czurko A, Mamiya A, Buzsáki G (1999) Oscillatory coupling of hippocampal pyramidal cells and interneurons in the behaving Rat. Journal of Neuroscience 19:274–287.

Davidson TJ, Kloosterman F, Wilson MA (2009) Hippocampal replay of extended experience. Neuron 63:497–507.

Di Prisco GV, Vertes RP (2006) Excitatory actions of the ventral midline thalamus (rhomboid/reuniens) on the medial prefrontal cortex in the rat. Synapse 60:45–55.

Dolleman-Van Der Weel MJ, Witter MP (1996) Projections from the nucleus reuniens thalami to the entorhinal cortex, hippocampal field CA1, and the subiculum in the rat arise from different populations of neurons. The Journal of comparative neurology 364:637–650.

Dolleman-Van der Weel MJ, Witter MP (2000) Nucleus reuniens thalami innervates gamma aminobutyric acid positive cells in hippocampal field CA1 of the rat. Neuroscience letters 278:145–148.

Dolleman-Van der Weel MJ, Lopes da Silva FH, Witter MP (1997) Nucleus reuniens thalami modulates activity in hippocampal field CA1 through excitatory and inhibitory mechanisms. Journal of neuroscience 17:5640–5650.

Dolleman-van der Weel MJ, Griffin AL, Ito HT, Shapiro ML, Witter MP, Vertes RP, Allen TA (2019) The nucleus reuniens of the thalamus sits at the nexus of a hippocampus and medial prefrontal cortex circuit enabling memory and behavior. Learn Mem 26:191–205.

Dragoi G, Buzsáki G (2006) Temporal encoding of place sequences by hippocampal cell assemblies. Neuron 50:145–157.

Dragoi G, Tonegawa S (2012) Preplay of future place cell sequences by hippocampal cellular assemblies. Nature 469:397–401.

Eichenbaum H (2017) Prefrontal-hippocampal interactions in episodic memory. Nature Rev Neurosci 18:547–558.

Euston DR, Tatsuno M, McNaughton BL (2007) Fast-forward playback of recent memory sequences in prefrontal cortex during sleep. Science 318:1147–1150.

Ferezou I, Bolea S, Petersen CC (2006) Visualizing the cortical representation of whisker touch: voltage-sensitive dye imaging in freely moving mice. Neuron 50:617–629.

Ferraris M, Ghestem A, Vicente AF, Nallet-Khosrofian L, Bernard C, Quilichini PP (2018) The Nucleus Reuniens Controls Long-Range Hippocampo-Prefrontal Gamma Synchronization during Slow Oscillations. J Neurosci 38:3026–3038.

Foster DJ (2017) Replay Comes of Age. Annu Rev Neurosci 40:581–602.

Foster DJ, Wilson MA (2006) Reverse replay of behavioural sequences in hippocampal place cells during the awake state. Nature 440:680–683.

Frankland PW, Bontempi B (2005) The organization of recent and remote memories. Nature Reviews Neuroscience 6:119–130.

Harris KD, Henze DA, Csicsvari J, Hirase H, Buzsáki G (2000) Accuracy of tetrode spike separation as determined by simultaneous intracellular and extracellular measurements. Journal of Neurophysiology 84:401–414.

Harris KD, Csicsvari J, Hirase H, Dragoi G, Buzsáki G (2003) Organization of cell assemblies in the hippocampus. Nature 424:552–556.

Hazan L, Zugaro M, Buzsáki G (2006) Klusters, NeuroScope, NDManager: a free software suite for neurophysiological data processing and visualization. J Neurosci Methods 155:207–216.

Herkenham M (1978) The connections of the nucleus reuniens thalami: evidence for a direct thalamo-hippocampal pathway in the rat. The Journal of comparative neurology 177:589–610.

Hoover WB, Vertes RP (2012) Collateral projections from nucleus reuniens of thalamus to hippocampus and medial prefrontal cortex in the rat: a single and double retrograde fluorescent labeling study. Brain structure & function 217:191–209.

Hutt A (2011) Sleep and Anesthesia; Neural correlates in Theory and experiment: Springer.

Isomura Y, Sirota A, Ozen S, Montgomery S, Mizuseki K, Henze DA, Buzsáki G (2006) Integration and segregation of activity in entorhinal-hippocampal subregions by neocortical slow oscillations. Neuron 52:871–882.

Ito HT, Zhang S-j, Witter MP, Moser EI, Moser M-b (2015) A prefrontal–thalamo– hippocampal circuit for goal-directed spatial navigation. Nature 522:50–55.

Jankowski MM, Islam MN, Wright NF, Vann SD, Erichsen JT, Aggleton JP, O’Mara SM (2014) Nucleus reuniens of the thalamus contains head direction cells. eLife 3:e03075–e03075.

Jankowski MM, Passecker J, Islam MN, Vann S, Erichsen JT, Aggleton JP, O’Mara SM (2015) Evidence for spatially-responsive neurons in the rostral thalamus. Frontiers in Behav Neurosci 9:256–256.

Ji D, Wilson MA (2007) Coordinated memory replay in the visual cortex and hippocampus during sleep. Nature Neuroscience 10:100–107.

Jones EG (1985) The Thalamus. Plenum Press, New York 1985: Springer.

Kenet T, Bibitchkov D, Tsodyks M, Grinvald A, Arieli A (2003) Spontaneously emerging cortical representations of visual attributes. Nature 425:954–956.

Khodagholy D, Gelinas JN, Buzsaki G (2017) Learning-enhanced coupling between ripple oscillations in association cortices and hippocampus. Science 358:369–372.

Kitamura T, Ogawa SK, Roy DS, Okuyama T, Morrissey MD, Smith LM, Redondo RL, Tonegawa S (2017) Engrams and circuits crucial for systems consolidation of a memory. Science 356:73–78.

Lansink CS, Goltstein PM, Lankelma JV, McNaughton BL, Pennartz CM (2009) Hippocampus leads ventral striatum in replay of place-reward information. PLoS Biol 7:e1000173–e1000173.

Latchoumane C-FV, Ngo H-VV, Born J, Shin H-S (2017) Thalamic Spindles Promote Memory Formation during Sleep through Triple Phase-Locking of Cortical, Thalamic, and Hippocampal Rhythms. Neuron 95:424–435.

Lee AK, Wilson MA (2002) Memory of sequential experience in the hippocampus during slow wave sleep. Neuron 36:1183–1194.

Liu K, Sibille J, Dragoi G (2019) Preconfigured patterns are the primary driver of offline multi-neuronal sequence replay. Hippocampus.

Loureiro M, Cholvin T, Lopez J, Merienne N, Latreche A, Cosquer B, Geiger K, Kelche C, Cassel J-C, Pereira de Vasconcelos A (2012) The ventral midline thalamus (reuniens and rhomboid nuclei) contributes to the persistence of spatial memory in rats. Journal of Neuroscience 32:9947–9959.

Luczak A, Maclean JN (2012) Default activity patterns at the neocortical microcircuit level. Frontiers in integrative neuroscience 6:30–30.

Luczak A, McNaughton BL, Harris KD (2015) Packet-based communication in the cortex. Nature Reviews Neuroscience 16:745–755.

Luczak A, Bartho P, Marguet SL, Buzsáki G, Harris KD (2007) Sequential structure of neocortical spontaneous activity in vivo. PNAS 104:347–352.

Maass W (2016) Searching for principles of brain computation. Current Opinion in Behavioral Sciences 11:81–92.

MacLean JN, Watson BO, Aaron GB, Yuste R (2005) Internal dynamics determine the cortical response to thalamic stimulation. Neuron 48:811–823.

Maingret N, Girardeau G, Todorova R, Goutierre M, Zugaro M (2016) Hippocampo-cortical coupling mediates memory consolidation during sleep. Nature neuroscience 19:959–964.

Malvache A, Reichinnek S, Villette V, Haimerl C, Cossart R (2016) Awake hippocampal reactivations project onto orthogonal neuronal assemblies. Science 353:1280–1283.

Manouze H, Ghestem A, Poillerat V, Bennis M, Ba-M’hamed S, Benoliel JJ, Becker C, Bernard C (2019) Effects of Single Cage Housing on Stress, Cognitive, and Seizure Parameters in the Rat and Mouse Pilocarpine Models of Epilepsy. eNeuro 6.

Marre O, Yger P, Davison AP, Frégnac Y (2009) Reliable Recall of Spontaneous Activity Patterns in Cortical Networks. Journal of Neuroscience 29:14596–14606.

Nadasdy Z, Hirase H, Czurko A, Csicsvari J, Buzsáki G (1999) Replay and time compression of recurring spike sequences in the hippocampus. Journal of Neuroscience 19:9497–9507.

Nádasdy Z (2000) Spike sequences and their consequences. Journal of physiology, Paris 94:505–524.

Pastalkova E, Itskov V, Amarasingham A, Buzsaki G (2008) Internally generated cell assembly sequences in the rat hippocampus. Science 321:1322–1327.

Pereira de Vasconcelos A, Cassel J-C (2015) The nonspecific thalamus: A place in a wedding bed for making memories last? Neurosci & biobehav rev 54:175–196.

Peyrache A, Khamassi M, Benchenane K, Wiener SI, Battaglia FP (2009) Replay of rule-learning related neural patterns in the prefrontal cortex during sleep. Nature neuroscience 12:919–926.

Pfeiffer BE (2017) The Content of Hippocampal “Replay”. Hippocampus:1–13.

Preston AR, Eichenbaum H (2013) Interplay of hippocampus and prefrontal cortex in memory. Current biology: CB 23:R764–773.

Quilichini P, Sirota A, Buzsáki G (2010) Intrinsic circuit organization and theta-gamma oscillation dynamics in the entorhinal cortex of the rat. J Neurosci 30:11128–11142.

Sanchez-Vives MV, Mattia M (2014) Slow wave activity as the default mode of the cerebral cortex. Archives Italiennes de Biologie 152:147–155.

Siapas AG, Wilson MA (1998) Coordinated interactions between hippocampal ripples and cortical spindles during slow-wave sleep. Neuron 21:1123–1128.

Sirota A, Buzsáki G (2005) Interaction between neocortical and hippocampal networks via slow oscillations. Thalamus & related systems 3:245–259.

Sirota A, Csicsvari J, Buhl D, Buzsáki G (2003) Communication between neocortex and hippocampus during sleep in rodents. PNAS 100:2065–2069.

Skaggs WE, McNaughton BL (1996) Replay of neuronal firing sequences in rat hippocampus during sleep following spatial experience. Science 271:1870–1873.

Staresina BP, Ole Bergmann T, Bonnefond M, van der Meij R, Jensen O, Deuker L, Elger CE, Axmacher N, Fell J (2015) Hierarchical nesting of slow oscillations, spindles and ripples in the human hippocampus during sleep. Nature Neuroscience 18:1679–1686.

Tonegawa S, Morrissey MD, Kitamura T (2018) The role of engram cells in the systems consolidation of memory. Nat Rev Neurosci 19:485–498.

Tung A, Mendelson WB (2004) Anesthesia and sleep. Sleep Med Rev 8:213–225.

Van der Werf YD, Witter MP, Groenewegen HJ (2002) The intralaminar and midline nuclei of the thalamus. Anatomical and functional evidence for participation in processes of arousal and awareness. Brain Res Brain Res Rev 39:107–140.

Varela C, Kumar S, Yang JY, Wilson MA (2014) Anatomical substrates for direct interactions between hippocampus, medial prefrontal cortex, and the thalamic nucleus reuniens. Brain Struct Funct 219:911–929.

Vertes RP (2004) Differential projections of the infralimbic and prelimbic cortex in the rat. Synapse 51:32–58.

Vertes RP (2006) Interactions among the medial prefrontal cortex, hippocampus and midline thalamus in emotional and cognitive processing in the rat. Neuroscience 142:1–20.

Villette V, Malvache A, Tressard T, Dupuy N, Villette V, Malvache A, Tressard T, Dupuy N, Cossart R (2015) Internally Recurring Hippocampal Sequences as a Population Template of Spatiotemporal Information Article Internally Recurring Hippocampal Sequences as a Population Template of Spatiotemporal Information. Neuron 88:357–366.

Walsh DA, Brown JT, Randall AD (2017) In vitro characterization of cell-level neurophysiological diversity in the rostral nucleus reuniens of adult mice. J Physiol 595:3549–3572.

